# A functional-structural connectivity metric detects ipsilateral connections with distinct functional specialisation in each hemisphere

**DOI:** 10.1101/2020.12.03.410902

**Authors:** Oren Civier, Marion Sourty, Fernando Calamante

## Abstract

We introduce a connectomics metric that integrates information on structural connectivity (SC) from diffusion MRI tractography and functional connectivity (FC) from resting-state functional MRI, at individual subject level. The metric is based on the ability of SC to broadly predict FC using a simple linear predictive model; for each connection in the brain, the metric quantifies the deviation from that model. For the metric to capture underlying physiological properties, we minimise systematic measurement errors and processing biases in both SC and FC, and address several challenges with the joint analysis. This also includes a data-driven normalisation approach. The combined metric may provide new information by indirectly assessing white matter structural properties that cannot be inferred from diffusion MRI alone, and/or complex interregional neural interactions that cannot be inferred from functional MRI alone. To demonstrate the utility of the metric, we used young adult data from the Human Connectome Project to examine all bilateral pairs of ipsilateral connections, i.e. each left-hemisphere connection in the brain was paired with its right-hemisphere homologue. We detected a minority of bilateral pairs where the metric value is significantly different across hemispheres, which we suggest reflects cases of ipsilateral connections that have distinct functional specialisation in each hemisphere. The pairs with significant effects spanned all cortical lobes, and also included several cortico-subcortical connections. Our findings highlight the potential in a joint analysis of structural and functional measures of connectivity, both for clinical applications and to help in the interpretation of results from standard functional connectivity analysis.

**Significance Statement:** Based on the notion that structure predicts function, the scientific community sought to demonstrate that structural information on fibre bundles that connect brain regions is sufficient to estimate the strength of interregional interactions. However, an accurate prediction using MRI has proved elusive. This paper posits that the failure to predict function from structure originates from limitations in measurement or interpretation of either diffusion MRI (to assess fibre bundles), fMRI (to assess functional interactions), or both. We show that these limitations can be nevertheless beneficial, as the extent of divergence between the two modalities may reflect hard-to-measure properties of interregional connections, such as their functional role in the brain. This provides many insights, including into the division of labour between hemispheres.

## Introduction

System-level brain organization can be described based both on structural connectivity and functional connectivity. Logically, structural connectivity should give rise to functional connectivity, and indeed, many previous studies have shown that measured properties of structural connections relate to those of functional connections (1). Despite these advances, there have been few attempts to quantitatively combine these two types of measurements for clinical research, with most studies still running separate analyses for structural and functional connectivity data (e.g., ref. 2). Here we introduce a metric that quantitatively combines information from both structural and functional connectivity, to reveal brain organisation properties not directly measured by either of them separately, such as hemispheric differences in functional specialization.

The metric we introduce is based on the ability of a connection’s structural properties to *broadly* predict its functional properties (3–5). To estimate this relationship, most previous studies measured the properties of structural connections using tractography algorithms applied to diffusion MRI (dMRI) data and quantified each connection’s ‘strength’ using a single value (which we will denote SC, for ‘structural connectivity’). Similarly, the properties of functional connections are usually estimated by applying time course correlation analysis to functional MRI (fMRI) data, and each connection’s ‘strength’ is again quantified by a single value (which we will denote FC, for ‘functional connectivity’). Early connectivity studies hypothesized (which turned out to be correct) a positive relationship between the SC and FC of interregional connections, as more axons/larger axon diameter/thicker myelin sheaths (structural properties captured by dMRI tractography) should lead to greater functional interactions between regions, hence, increased time course correlation (6). This is assuming that most interregional connections in the brain are excitatory rather than inhibitory, such that the activity levels of two inter-connected regions tend to fluctuate in the same direction. Unfortunately, however, the prediction of FC by SC has been found to be limited (7, 8). Indeed, from the many studies that tackled the problem, SC could usually explain no more than ~50% of the variance in FC for parcellations with tens of regions (3–5), or no more than ~30% of variance for parcellations with hundreds of regions (3, 4).

Based on theoretical considerations that question the ability of dMRI tractography to capture the full characteristics of white matter (8, 9), or of a simple correlation-based measure to capture complex neural interactions (8, 10, 11), we here part our ways from previous investigations: instead of trying to account for the variation in FC still unexplained by SC, we propose to rather exploit this mismatch between the two modalities. For each connection, the metric that we introduce aims therefore to measure the amount of mismatch between two values: empirical FC, and the FC predicted from SC by a simple linear regression model (the model that is commonly used in previous studies, see ref. 3, 4, 12). We will denote this mismatch the *FC-SC mismatch metric*, and under the assumption that the mismatch has, at least in part, a physiological basis, we will use it to indirectly quantify certain physiological properties not accessible when analysing SC and FC separately.

The FC-SC mismatch metric builds on the notion that dMRI tractography and time-course fMRI correlations do perform well, respectively, in capturing the strength of SC in connections with ‘*common’-structure* fibre bundles (‘common’ in the sense that the axons of the connection do not have special features such as extensive branching), and the strength of FC in connections with ‘*simple’* neural interactions (‘simple’ in the sense that the neural interactions consist of one region driving the activity in the other one, without the connection being involved in more complex neural computations such as those required for the implementation of Boolean operations).

Because the strength of SC is usually assumed to be a direct driver for the strength of FC, the relationship between SC and FC in these typical connections should be well-characterized then by a simple linear regression model (3). If all connections in the brain were of this pattern (and assuming no systematic measurement errors or processing biases), the predictive regression model between SC and FC would fit perfectly, with a 0 mismatch value (i.e. FC-SC mismatch metric equals 0) for each of the connections. However, some connections diverge from this pattern by having ‘special’ white matter structural properties, e.g., extensive branching in the form of axon collaterals, that are not well-captured by dMRI tractography (either not captured at all, or measured incorrectly). Yet, other connections, or sometimes even the same connections, may diverge from this pattern by being involved in complex neural interactions, which, similarly to the dMRI case, are not well-captured by time-course correlations of the fMRI measurement; for example, an “OR” Boolean operation. These measurement limitations/inaccuracies may cause the SC or/and FC of some connections to deviate from the regression line, thus giving rise to FC-SC mismatches.

A possible application of the FC-SC mismatch metric could be in testing whether each connection’s metric is significantly different from 0, i.e. testing if the connection has ‘special’ white matter structural properties and/or complex neural interactions. However, this approach has two main caveats. First, the range of neural interactions and white matter structural properties in the brain is enormous, and since all of them contribute to the simple linear regression model (including all the cases where the measure of SC or FC is inaccurate), the model is unlikely to represent the ‘pure’ relationship between SC and FC described above. Instead, the model represents some ‘average’ relationship between SC and FC, which is still broadly valid, but not accurate enough for deciding whether a specific connection is atypical. The second caveat is that calculating SC and FC is affected by systematic measurement errors and processing biases (9, 13). These errors and biases may be somewhat minimised, but unfortunately, some of them cannot be accounted for effectively using existing methods (e.g., errors introduced into tractography by white matter bottlenecks, see ref. 14). With these caveats in mind, we suggest to limit the scope of the FC-SC mismatch metric, at least for the illustrative application shown here: rather than comparing each connection’s metric value to zero, we will use the metric to compare different connections, and, specifically, for comparison between connections that share equivalent systematic measurement errors and processing biases (thus cancelling out these sources of error).

To illustrate an important application of the method, we take advantage of the fact that systematic errors and processing biases in both SC and FC are likely comparable bilaterally, and consider the case of comparing pairs of intra-hemispheric connections: a connection in the left hemisphere, and its right-hemisphere homologue. We will refer to each of these pairs as a *bilateral pair* and evaluate the two associated FC-SC mismatch metric values, for left and right *homologues*. Notice that we will not be interested in the metric values per-se, but rather in whether they are the same or different across the hemispheres. By directly comparing left and right FC-SC mismatches within each bilateral pair, we will detect cases where complex neural interactions and/or white matter structural properties (those not well-captured by dMRI tractography) likely vary across the hemispheres. This could point to bilateral pairs with distinct functional specialisation in each hemisphere, i.e. each homologue fulfils a completely different role in the brain.

### Metric

Tractography biases are arguably the largest source for systematic errors that might affect metrics that rely on structural connectivity. To lessen this problem, our proposed metric uses the state-of-the-art quantitative tractography method, SIFT2 (15), which is specifically designed to reduce tracking biases (this can be substituted with comparable methods, e.g., ref. 16–19). Besides using advanced quantitative tracking, two other main challenges of defining an FC-SC mismatch metric were addressed (see Fig. 1). The first challenge is the long tail that is present in the SC distribution (i.e. the SC values for all connections within an individual’s brain) but not in the FC distribution, indicating that only in the former there is a small portion of connections that are much stronger than the rest (20). When applying a linear regression model during metric calculation, this discrepancy tends to invalidate the assumption of homoscedasticity of the residual errors. We solve it using a data-driven transformation based on a power-law, which makes the distribution of SC similar to that of FC. The second challenge is that FC quantifies connectivity strength based on time-correlated activity which is driven by the single *direct* and many *indirect* structural links (or simply, links) between each pair of regions; in contrast, SC quantifies connectivity by measuring the actual fibre bundle that connects each pair of regions, and this only accounts for the *direct* link between them. There were attempts at incorporating some information on indirect links into SC after all (4), but they unfortunately fell short of resolving this intrinsic inter-modal discrepancy. Conversely, our approach aims to evade the problem rather than solving it. Treating the regions and connections in the brain as a graph (a ‘connectome’, ref. 21), we apply a local graph-theoretical metric to identify connections where *indirect* links are major contributors to FC. We mark these connections as unsuitable for calculation of the FC-SC mismatch metric, and exclude them from further analysis.

**Figure 1.**
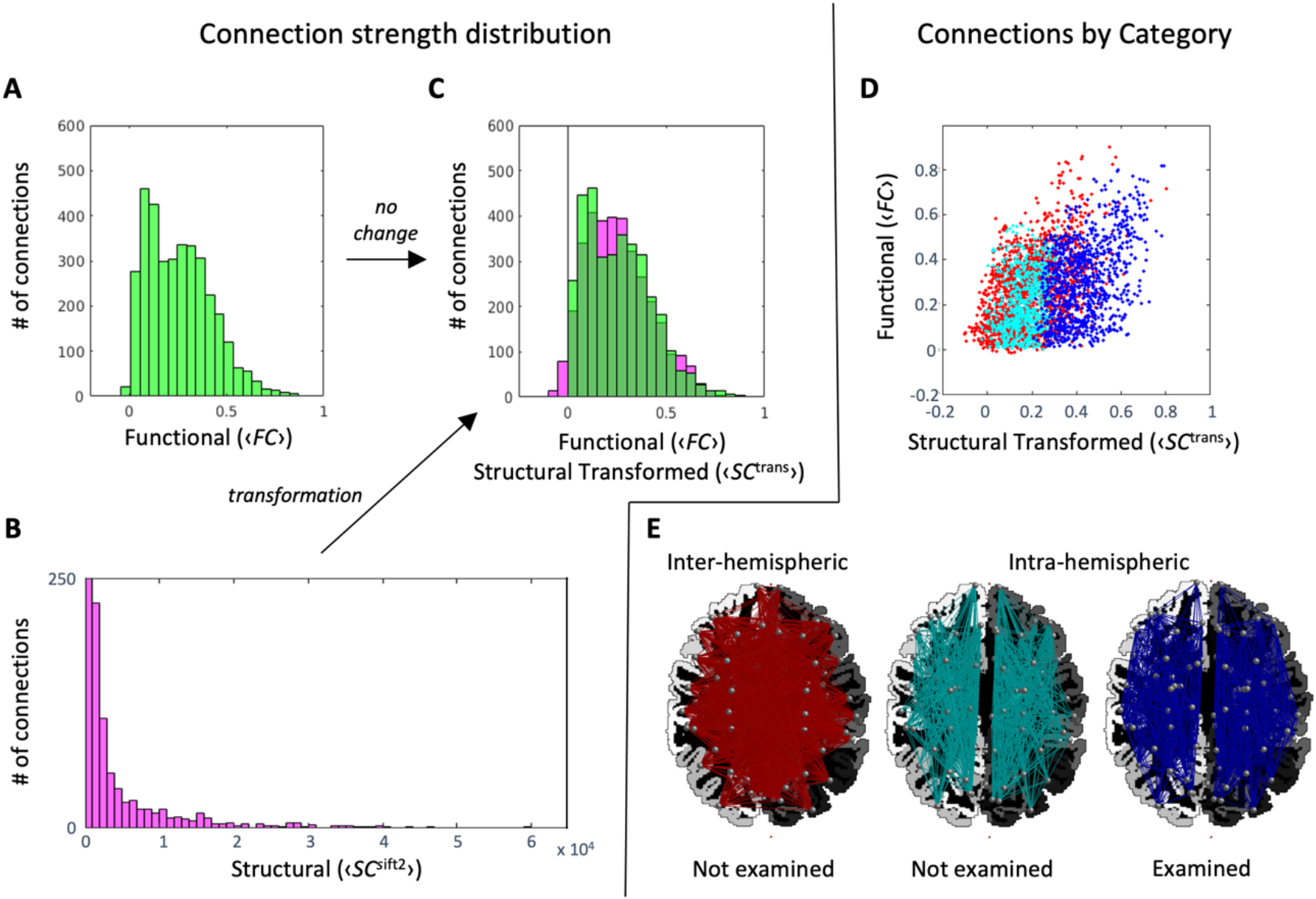
Processing of functional connectivity and structural connectivity data. All data shown are group-averaged. SC transformation by histogram fitting to FC: (*A*) Distribution of ‹*FC*›. (*B*) Distribution of ‹*SC*^sift2^*›*. For visualisation, y-axis is cut off at 250 (excluding most of the leftmost bar, which extends to 2760 connections), and x-axis at 6,500 (excluding the four strongest connections in the distribution). (*C*) Distribution of ‹*SC*^trans^› = −0.3789 + 0.4114*(‹*SC*^sift2^›)0.0926 overlaid on the distribution of ‹*FC*› (see *Materials and Methods* for details). Green - ‹*FC*›; Magenta - ‹*SC*^trans^›; Dark green - their overlap. Exclusion of connections from the analysis: (*D*) Scatter plot of ‹*FC*› against ‹SCtrans›. Red, excluded – the 1764 inter-hemispheric connections (including 42 homotopic connections); Cyan, excluded – the 770 intra-hemispheric connections where there is at least one indirect path shorter than the direct path (according to graph-theoretical definitions, see *Materials and Methods*); Blue, preserved – the 952 intra-hemispheric connections where the direct path is the shortest. (*E*) The connections in *D*, plotted on a representative brain. ‹ › indicate group-averaged.

After the above two steps and the additional exclusion of inter-hemispheric connections (further details and equations available in *Materials and Methods*), a valid and biologically-meaningful characterization of the relationship between SC and FC using a linear regression model finally becomes feasible. The fit of the simple linear regression model *f*_*k*_ is performed for each individual subject *k* separately in order to capture the (poorly understood) inter-individual variation in the range (scale) and mean of the FC distribution (13).

### Definition of the FC-SC *mismatch* metric

The metric *mFCSC*_*uv,k*_ is defined as:

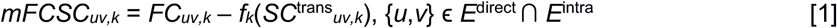

where *{u, v}* is a connection between regions *u≠v*, *k* is subject index, *FC*_*uv,k*_ is functional connectivity for connection *{u,v}* of subject *k*, *SC*^trans^_*uv,k*_ is transformed SC for the same connection (Eq. 3), and *f*_*k*_ is the linear model that fits FC to transformed SC. *E*^direct^ is the set of connections where the direct link between the regions is the major contributor to FC (Eq. 4), and *E*^*intra*^ is the set of all intra-hemispheric connections in the brain. The metric *mFCSC* is the residual of the simple linear regression of *FC*_*uv,k*_ against *SC*^tran*s*^*uv,k*. Geometrically, it is the y-axis difference of *FC*_*uv,k*_ from the regression line for subject *k* (see Fig. 2).

**Figure 2.**
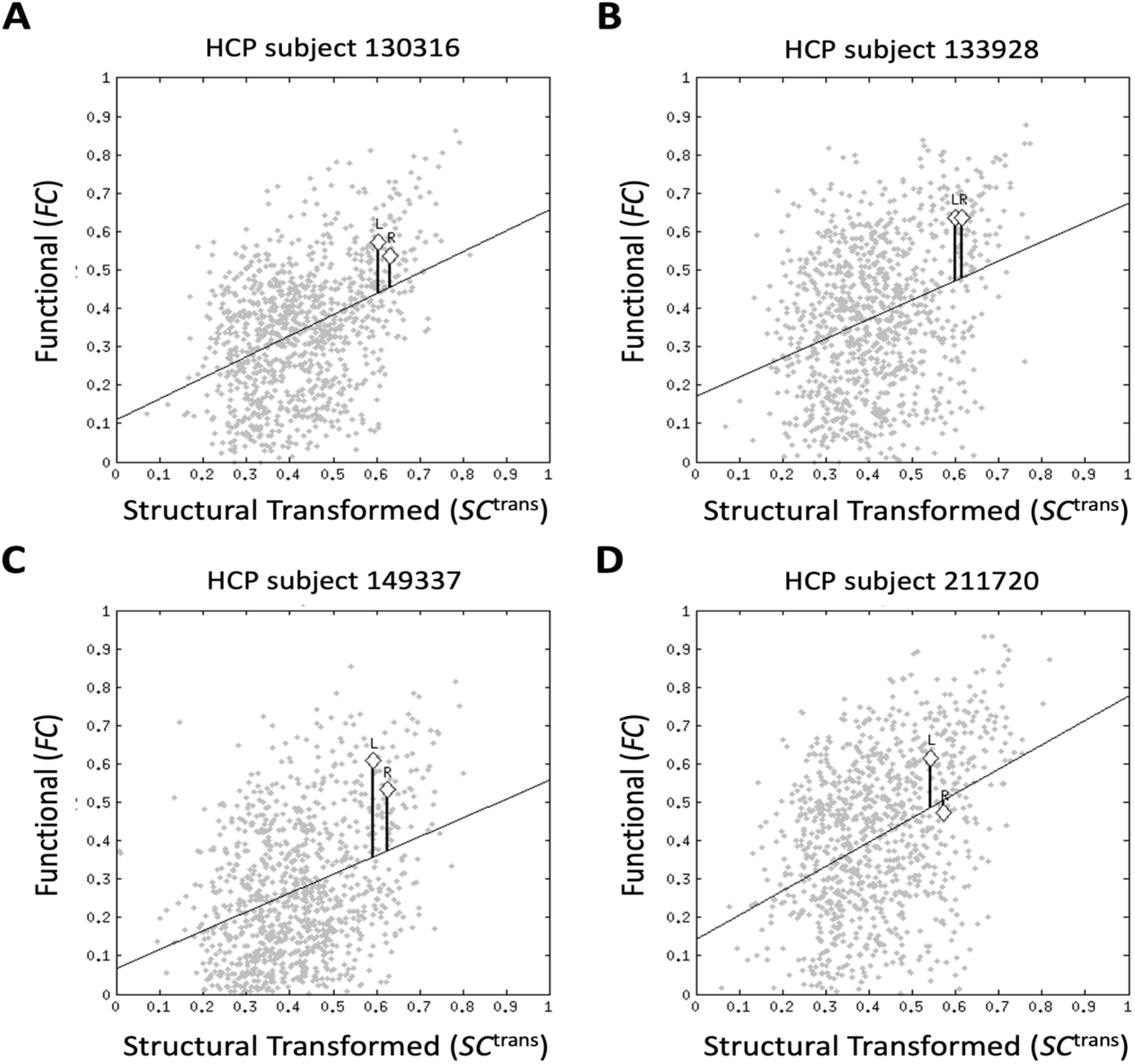
Scatterplots of functional connectivity against transformed structural connectivity (grey points) and corresponding regression lines for 4 representative subjects from the Human Connectome Project dataset. To give an example of a bilateral pair that had significantly different FC-SC mismatch in each hemisphere, the data points for the left and right superior frontal*-*pars opercularis connections (frontal aslant tracts, or FATs) are indicated by diamonds. The y-axis distance of each data point from the regression line is equal to its FC-SC mismatch metric value, i.e. *mFCSC*_*uv,k*_. The distances are indicated by vertical lines. HCP = Human Connectome Project. L = left homologue. R = right homologue.

## Results

For this study, we processed dMRI and fMRI data from healthy young adults to obtain individual SC and FC connectomes using the 84 regions of the Desikan-Killiany atlas (22). We first estimated the parameters of the power-law function, applied it to the group-averaged SC connectome, and excluded all inter-hemispheric as well as some intra-hemispheric connections (Fig. 1; power-law parameters are *a* = −0.3789, *b* = 0.4114, and *c* = 0.0926; out of 1,722 intra-hemispheric connections, 770 were excluded). The coefficient of the Pearson correlation between the resulting transformed group-averaged SC (or ‹*SC*^trans^›), and the group-averaged FC (or ‹*FC*›), was *r* = 0.41. We then applied the power-law function to the individual SC connectomes, and fit a simple linear regression model between the FC and the transformed SC of each subject (Fig. 2). These pairs of structural and functional connectomes, as well as their corresponding linear models, were used in all subsequent analyses.

### Bilateral pairs with different FC-SC mismatch metric values

Let {*u*,*v*}[L] ϵ *E*^direct^, *u*≠*v* be an intra-hemispheric connection on the left hemisphere, and {*u*,*v*}[R] ϵ *E*^direc*t*^, the homologous connection on the right; together, they are a bilateral pair. In our cohort of 50 individuals there were, therefore, 50 instances of each such pair, which allowed using a paired two-sample t-test to test the hypothesis that *mFCSC*_*uv*[L]_ ≠ *mFCSC*_*uv*[R]_. The 41 bilateral pairs where this difference was significant (*p* < 0.00006; Bonferroni corrected for 861 bilateral pairs, with alpha set to 0.05) are given in Fig. 3 and Table S1. As we corrected for multiple comparisons across all 861 bilateral pairs in the brain, this result is not dependent on the number of connections excluded earlier in the analysis (which may vary between different cohorts).

**Figure 3.**
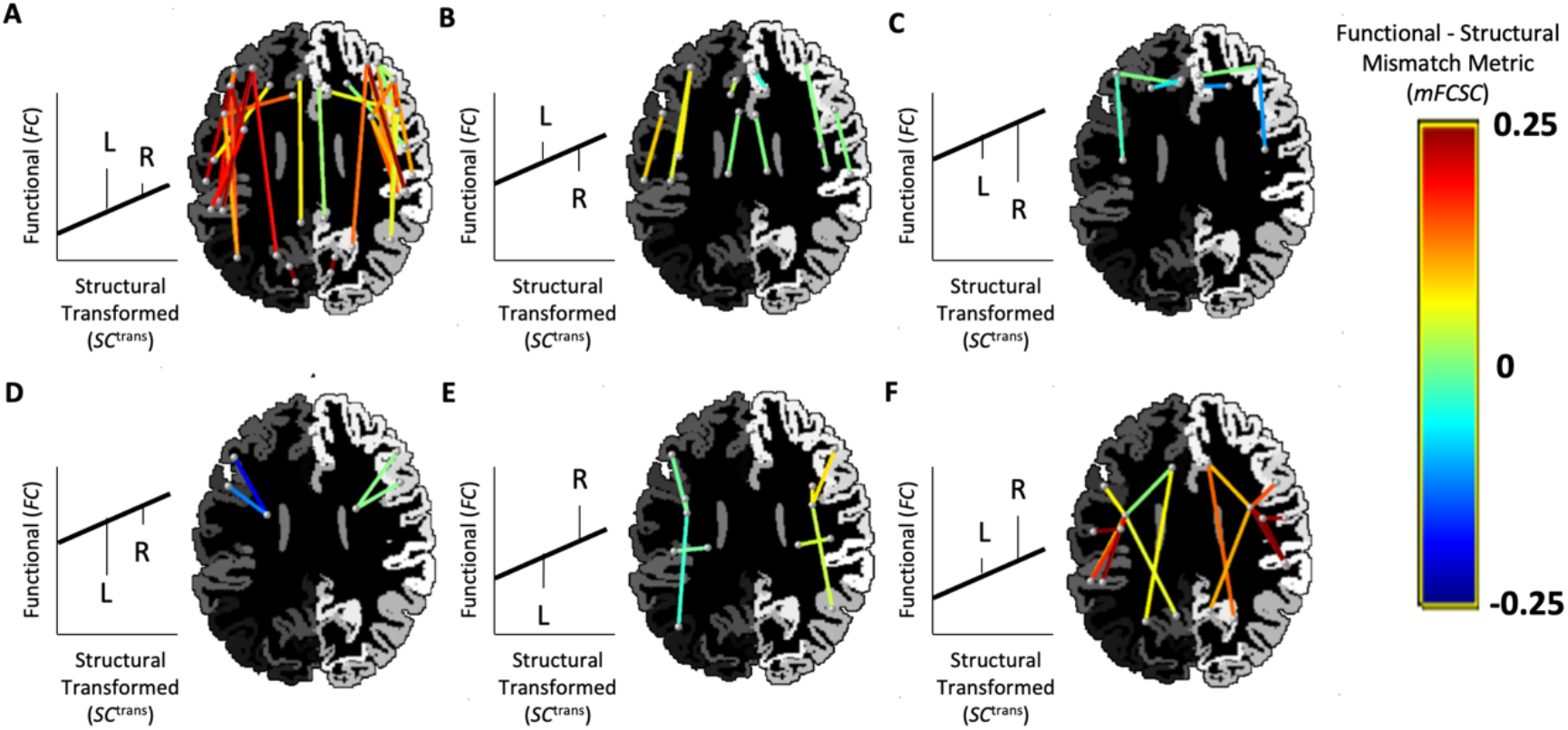
Bilateral pairs (of intra-hemispheric connections) that had significant difference between the FC-SC mismatch metric values in left and right homologues (*p* < 0.00006; Bonferroni corrected with alpha set to 0.05). Each pair is represented by two lines, for left- and right-hemisphere, respectively. The dots at the edges of the lines represent the connected regions, and are positioned at their centres. An axial slice of the parcellation for a representative brain serves as a background (but notice that the regions represented by the dots might be superior or inferior to that plane). The FS-SC mismatch metric value of each connection is given by the line colour. The bilateral pairs are divided into 6 panels (*A-F*) according to the relationship between the average values of *mFCSC* within each pair; this is schematically illustrated for an idealised subject on the left hand of each panel, and using the conventions of Fig. 2. For simplicity, structural connectivity (*SC*^trans^) in the illustrations is always stronger in the right homologue (the vertical line denoted by “R” is to the right of the “L” line), but in the actual data, there are also cases where this is reversed. L = left homologue. R = right homologue.

The bilateral pairs with significant effects spanned all cortical lobes, and also included several cortico-subcortical connections. The pairs may be categorised according to the relationship between the values of the mean left and right FC-SC mismatch metrics (‹*mFCSC*_[L]_› > ‹*mFCSC*_[R]_› or vice versa; Fig. 3, top and bottom rows) as well as their signs (different panels). It is notable that most pairs have positive mismatches in both left and right homologues (panels *A* and *F*), whereas only the minority of cases exhibit the opposite pattern (i.e. negative mismatches in both homologues, panels *C* and *D*). Fig. 2 gives individual examples for one of the pairs in Fig. 3*A* (superior frontal-pars opercularis).

### Bilateral pairs with FC asymmetry but comparable FC-SC mismatch metric values bilaterally

In a connectomics analysis, *functional connectivity asymmetry* refers to a situation where FC differs between the left and right homologues of a bilateral pair. Past studies that detected FC asymmetry during resting-state have often used the result to conclude on the direction of hemispheric dominance and its magnitude; i.e. which of the homologues contributes more to processing, and to what extent. However, this interpretation might sometimes be incorrect given that it is not always valid to assume that both homologues contribute to the same brain function. To find relevant bilateral pairs where this assumption is likely valid after all, we first detected all pairs with FC asymmetry (Fig. 4 *A* and *D*), and then excluded from them relevant bilateral pairs with *mFCSC*_*uv*[L]_ ≠ *mFCSC*_*uv*[R]_. For clarity, this was done separately for FC asymmetries towards the left and towards the right (Fig. 4 *B* and *E*). Several bilateral pairs survived this exclusion (Fig. 4 *C* and *F*, Table S2). These pairs exhibit both FC asymmetry and *mFCSC* mismatch values that are comparable bilaterally. There was only one such case with FC asymmetry towards the right (inferior parietal-rostral middle frontal; Fig. 4*F*), but it should be taken into account that there were relatively few bilateral pairs with rightward FC asymmetry to begin with (the initial functional connectivity analysis resulted in only 9 pairs with rightward FC asymmetry, Fig. 4*D*, compared with 25 pairs with leftward asymmetry, Fig. 4*A*).

**Figure 4.**
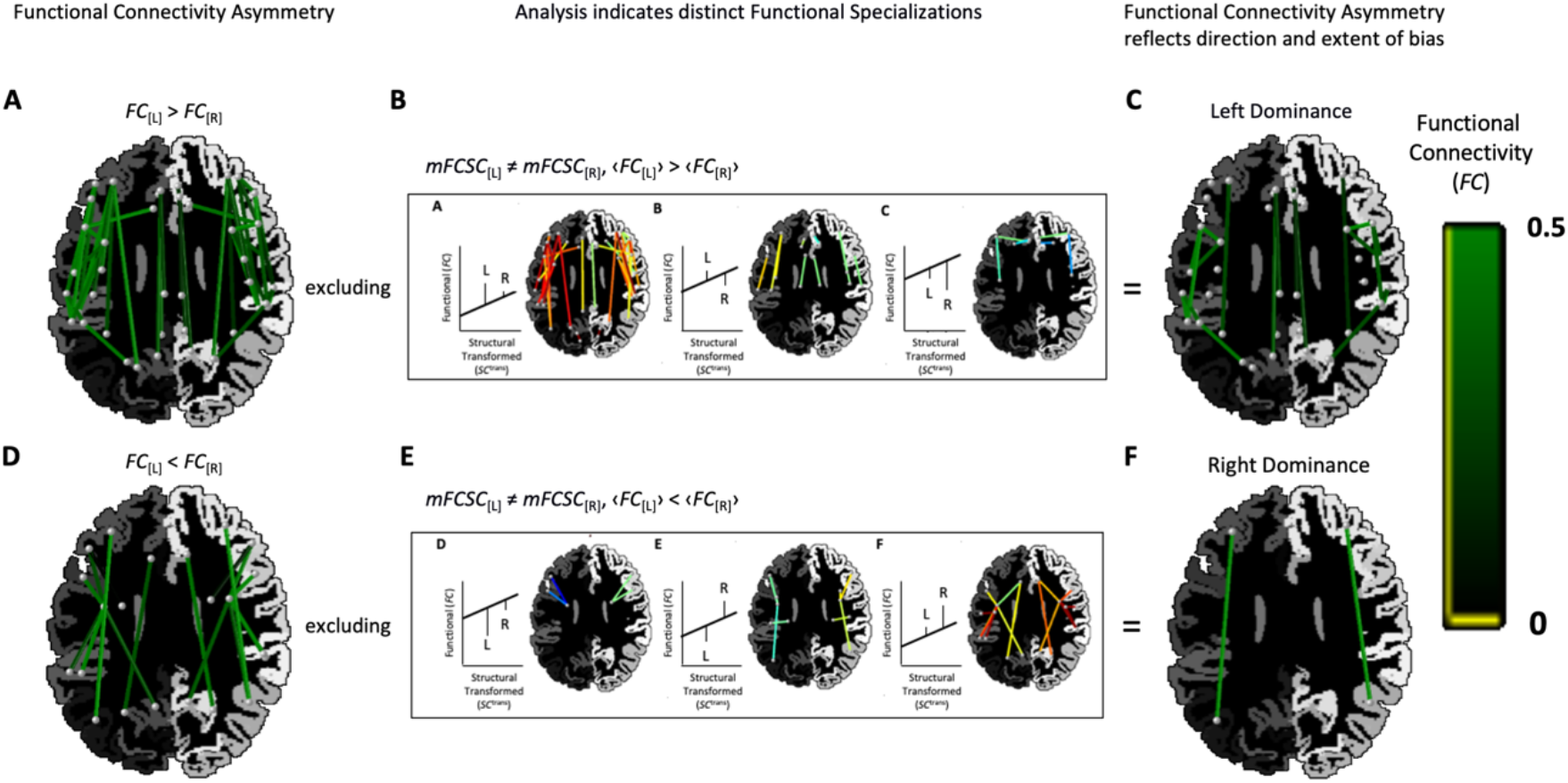
Detecting bilateral pairs where FC asymmetry is indicative of the direction and extent of functional hemispheric dominance. (*A*) Pairs that had left FC asymmetry, i.e. *FC*_[L]_ > *FC*_[R]_ (*p* < 0.00006; Bonferroni corrected with alpha set to 0.05), with colour intensity indicating FC strength, and other conventions as in Fig. 3, (*B*) excluding connections with *mFCSC*_[L]_ *≠ mFCSC*_[R]_ (connections to exclude correspond to the top row of Fig. 3, which is reproduced within the rectangle), (*C*) the subset of the pairs in panel *A* where left and right homologues probably share the same brain function (as *mFCSC* is not different between hemispheres). In these pairs, FC asymmetry is a good indicator for the direction and magnitude of functional hemispheric dominance, (*D*-*F*) Same conventions as panels *A*-*C*, but for the analysis of pairs that had right FC asymmetry.

## Discussion

We introduced a metric, FC-SC mismatch (*mFCSC*), which can be used to characterise the mismatch between the structural and functional connectivity of brain connections. To demonstrate its utility, we investigated in which bilateral pairs of intra-hemispheric connections the FC-SC mismatch is different between the two hemispheres. Importantly, the use of a connectome-based metric, which takes into account the relationship between SC and FC across all connections in the brain, allowed us to perform a highly sensitive analysis. We could thus run a whole-brain, yet well powered, analysis, indeed detecting multiple significant hemispheric differences.

The detected hemispheric differences may reveal aspects of brain organisation that cannot be investigated using only SC or FC in isolation. We suggest that this includes the identification of bilateral pairs where each homologue has distinct functional specialisation. To argue in favour of this interpretation, we first discuss the simplest case of bilateral pairs with no FC-SC mismatch at either hemisphere, and only then progress to the more sophisticated cases. For the discussion, we will use *SC*^trans^_*uv*[L]*,k*_ and *FC*_*uv*[L]*,k*_ (*SC*^trans^_*uv*[R],*k*_ and *FC*_*uv*[R],*k*_) for structural and functional connectivity of the left (right) homologue.

### Bilateral pairs without FC-SC mismatches

When *mFCSC*_*uv*[L],*k*_ ≈ *mFCSC*_*uv*[R],*k*_ ≈ 0 in an individual subject, then in both hemispheres, the relationship between *FC* and *SC*^trans^ is exactly the one predicted by the subject-specific linear model *f*_*k*_, indicating that both left and right homologues have only typical features (for this rather idealised example, we assume that the regression line simply represents the relationship between SC and FC in typical connections, and that there are no systematic measurement errors nor processing biases; cf. *Introduction*). Given that the hemispheres are largely symmetrical when it comes to the functional roles of brain regions (as reflected, for example, in the well-described resting-state functional networks being bilateral, see ref. 23, 24), we predict that the two homologues, being typical, agree with this general trend – hence, they share the same functional role in the brain. We cannot rule out that both data points fall on the regression line just by coincidence, but we consider it unlikely.

If the above equality occurs when the two data points have exactly the same *SC*^trans^ and *FC* (e.g., Fig. S1, circles), then this logically indicates that both homologues contribute equally to the specific brain function they share. In contrast, if the two data points differ in their SC (Fig. S1, pentagons), then the magnitude (but not the nature) of at least one property of the structural connections is likely different between the hemispheres (e.g., different number of axons). Due to the fact that the two data points are still on the regression line, this SC difference is obviously accompanied by a proportional relative difference in FC, which we consider to be the norm for typical connections (*Introduction*). This latter case may be interpreted as a situation when the two homologues share the same functional role, but with one of them contributing to a larger degree.

### Physiological basis for FC-SC mismatches

From this point on, we will focus on connections that do have a FC-SC mismatch, i.e. *FC* ≠ *f*_*k*_(*SC*^trans^). Assuming that the mismatch is not entirely due to systematic measurement errors or processing biases, a physiological basis may exist. A *positive* mismatch indicates that the empirical FC is larger than the FC predicted by SC. This often occurs when SC fails to capture special white matter structural properties because the underlying dMRI signal simply cannot measure them. For example, if in a given interregional fibre bundle, each axon is synapsing on an atypically high number of target neurons, then neurons in the source region will drive more neurons in the target region compared with the typical case. This may increase the time series correlation between the regions connected by the bundle. Because dMRI cannot measure these special microstructural properties, whereas fMRI can measure their consequence on the correlation strength, a positive FC-SC mismatch is thus to be expected. It is important to note that even if SC could reflect in its value the extensive synapsing, there is still no guarantee that this would correspond well enough with the increase in FC so as to zero out the FC-SC mismatch. Because special features such as extensive synapsing are atypical, it is still unknown whether the changes they introduce into SC and FC are proportional to each other, as is the case for the more typical connection features (*Introduction*). A *negative* mismatch, on the other hand, indicates that the empirical FC is smaller than the FC predicted by SC. This may occur when the connection is involved in complex neural interactions where some inputs modulate the others, such as an implementation of an AND operation. Inhibitory interregional axonal projections, though less frequent, can result in negative FC-SC mismatch as well; being mixed with excitatory projections that run between the same two regions, inhibitory projections would reduce the strength of the otherwise positive time series correlation. Another important source for both positive and negative mismatches is the organization of the grey matter regions themselves, and specifically, the existence of U-fibres. This is because the neural computation taking place internally in each of two connected brain regions, would also affect the nature of the neural interactions between them.

The different contributions detailed above are only a few examples out of many that can explain the physiological basis for the FC-SC mismatch. This unfortunately means that the *same* mismatch value may be generated by several *different* neural interactions and/or white matter structural properties (those not well-captured by dMRI tractography). It is impossible, therefore, to relate a specific mismatch value to a specific physiological factor, which limits this study to analysing the variation (but not the nature) of physiological factors across the brain.

### Pairs with comparable FC-SC mismatches in both homologues

If both left and right homologues have FC-SC mismatches different than 0, but comparable across the hemispheres, it is likely that both are affected by identical systematic measurement errors or processing biases. Alternatively, the source for the mismatches might be physiological, with the two homologues sharing the same atypical neural interactions and/or white matter structural properties (those not well-captured by dMRI tractography). While it is possible that the homologues do differ in these aspects, and by coincidence different physiological properties in each side gave rise to the same mismatch value, we consider it unlikely in light of the general tendency for symmetry discussed earlier in this section. The shared atypical features, in turn, suggest that both homologues also have a shared functional role; if each homologue has a different SC (and correspondingly, also a different FC; Fig. S1, squares), this may be considered a case of hemispheric dominance (see Fig. 4 *C* and *F*).

### Pairs with different FC-SC mismatch in left and right homologues

A case where the FC-SC mismatch is significantly different between the two homologues may indicate that both have complex neural interactions and/or atypical white matter structural properties (those not well-captured by dMRI tractography), but the exact nature of these traits is different in each hemisphere. It is also possible that only one of the homologues has atypical/complex properties, whereas both are affected by systematic measurement errors and processing biases (notice that the latter errors and biases are assumed to be identical across hemispheres, and thus cannot be the source of the hemispheric difference in *mFCSC*; see *Introduction*). Under the assumption that the brain is unlikely to use completely different mechanisms to fulfil the same functional role, this may indicate that the two homologues have different functional specialisations. Bilateral pairs with this configuration are at the focus of this study, and at the group level, we found 41 of them (Fig. 3).

To better understand the factors leading to different FC-SC mismatches in each hemisphere at the group-level, it is worth examining a single subject once again. The simple case occurs when *SC*^trans^_*uv,k*[L]_ ≈ *SC*^trans^_*uv,k*[R]_, but *mFCSC*_*uv,k*[L]_ ≠ *mFCSC*_*uv,k*[R]_ (Fig. S1, hexagons). This indicates that both homologues probably have the same number of axons / average axon diameter / level of myelination, all of these can be reasonably quantified by dMRI, whereas due to complex neural interactions in only one of the hemispheres, their FC values differ. Alternatively, it is possible that *atypical* white matter structural properties (that dMRI does not capture; hence, the equivalent SC values bilaterally) are the ones that are present in only one of the hemispheres, and the hemispheric difference in FC is due to the resulting increased/decreased neural interactions. The more complex case is characterised by differences in both *SC* and FC-SC mismatch, i.e. *SC*^trans^_*uv,k*[L]_ ≠ *SC*^trans^_*uv,k*[R]_ and *mFCSC*_*uv,k*[L]_ ≠ *mFCSC*_*uv,k*[R]_ (Fig. S1, diamonds). This indicates that the two homologues do not only have different functional specialisations, but each also requires a different level of neural resources (e.g., number of axons) -- in the example shown, the left homologue requires more resources, but due to a different functional specialisation compared with the right homologue, its FC is actually lower. Although not indicated for simplicity, the bilateral pairs shown in Fig. 3 can be either of the former simple case or the latter complex one.

### Comparison with neural-behavioural studies

The classical approach to detect functional specialisations is to examine the association of a selected connection with specific behaviour. If the conclusions from our analyses are valid, we expect them to agree with such studies. Although a thorough comparison is outside the scope of this study, there are several notable agreements. Based on our analysis (Fig. 2 and 3*A*, Table S1), we concluded that the bilateral superior frontal-pars opercularis connection (frontal aslant tract or FAT, see ref. 25) has distinct functional specialisation in each hemisphere; this is indeed consistent with neural-behavioural studies indicating a left FAT specialisation for speech actions, compared with a right FAT specialisation for general action control (mostly inhibitory control as required for stop-signal tasks) (26). Notice that although the superior frontal-pars triangularis connection form part of the FAT as well (26), it did not come up in our analysis. This is conceivably because the pars triangularis, in contrast with the pars opercularis, seems not to be associated with the speech production network in neither hemisphere (27). Another notable agreement concerns the nine insula connections that had significantly different FC-SC mismatch in each hemisphere (Fig. 3 *E* and *F*). Consistently, only the right insula is assumed to be a central node in a network involved in human body scheme representation, required for limb ownership and self-awareness of actions (28). Moreover, this network might be part of a larger right hemisphere network dominant for the perception of limb movement (29).

### Clinical implications

Owing to the results reported in this paper, clinicians would now be able to identify cases (e.g., using ref. 30) in which lesions affect bilateral pairs that have distinct functional specialisation in each hemisphere. It could be important, for example, in cases where only one of the two connections is damaged: because the contralateral homologue is specialised for another role, it is unlikely to compensate for the loss of function, and ipsilateral compensation is to be expected. This has consequences for both the strength of recovery -- possibly stronger in ipsilateral compensation (31) -- and its rate -- possibly slower in ipsilateral compensation (32) – which are two factors that must be considered when devising a rehabilitation plan. Lastly, the results of this paper will be especially valuable in cases where brain stimulation is used for treatment, and thus predicting the site of compensation becomes crucial (32).

Another application of the metric would suit longitudinal studies of multi-modal brain connectivity (e.g., ref. 33). In such an experimental design, we will not compare connections across the hemispheres, but rather, compare each connection to itself. Although participant repositioning in the scanner may pose a challenge for longitudinal studies of this sort, the related tractography biases have been shown to be effectively mitigated by SIFT (34). This application of the metric could be used, for example, to investigate whether changes in a connection’s FC from one time point to another are driven by plasticity/degeneration in direct links (*SC*^trans^ is changing with *FC*, and thus FC-SC mismatch is not changing over time) or indirect links (*SC*^trans^ is not changing with *FC*; FC-SC mismatch does change significantly over time). The results would inform studies on changes in structural-functional coupling during brain development and ageing (33, 35), and might be also useful in the evaluation of post-lesion plasticity (as long as fibre tracking through the lesioned region is feasible and valid). Lastly, this specific application takes into account indirect links, and thus, connections where they play a part (Fig. 1 *D* and *E)* should not be excluded from analysis anymore.

### Application for studying functional hemispheric dominances

A popular indicator for the direction and extent of *functional hemispheric dominance*, i.e. an intra-hemispheric connection that contributes to processing more than its contralateral homologue, is FC asymmetry (24). The weakness of this indicator, however, is that in bilateral pairs that exhibit a *distinct functional specialisation* in each hemisphere (Fig. 3), the FCs of the left and right homologues are independent of each other (as they may arise from completely different underlying neural interactions). In these cases, the value of FC asymmetry is arbitrary, and thus an unsuitable indicator. It is important to note that functional hemispheric dominance still exists in these pairs with distinct functional specialisations -- one brain function is dominant in the left hemisphere, and the other, in the right – the caveat is that to estimate the extent of left and right biases, measures other than resting-state fMRI based FC asymmetry must be used.

The analysis in Fig. 4 was aimed at detecting these cases where FC asymmetry is a reliable measure of functional hemispheric dominance after all. We first detected the bilateral pairs where FC asymmetry is significant to begin with, and after excluding bilateral pairs where the FC-SC mismatch metric is significantly different between hemispheres, we were only left with these pairs that have both significant FC asymmetry and similar functional roles bilaterally. In these bilateral pairs, FC asymmetry can be safely used to estimate both the direction and extent of hemispheric dominance.

### Limitations

The main limitation of the FC-SC mismatch metric is in the assumption that the non-linear model used to fit between the ‹*SC*^sift2^› and ‹*FC*› distributions at the group level, as well as the linear model used to fit between *SC*^trans^ and *FC* at subject-level, are suitable models. Based on the analyses presented here, the models do appear to provide a reasonable characterisation of the data. That said, the proposed metric can be generalised to explore the use of alternative, possibly more complex, models, which could potentially improve its accuracy. Many parameters that affect the FC-SC relationship are unknown; however, some parameters can be measured using imaging (e.g., distance between brain regions) or deduced from other sources (e.g., tracing studies indicating which connections are inhibitory), and could be incorporated into the models (11). It might be also possible to fit the distributions of *SC*^sift2^ and *FC* for each individual separately rather than at group-level. However, this would require assurances as to the stability of the model parameters fitted to the more atypical individuals.

Another limitation is that the intrinsic discrepancy between structural connectivity and functional connectivity (due to the existence of indirect links, see *Metric*) is evaded here by simply excluding the most affected connections from analysis. Moreover, the procedure uses a hard threshold, i.e. connections are excluded only if the indirect link is shorter than the direct link (according to graph-theoretical definition of path length), which is suboptimal. It would be preferable to use instead a more flexible exclusion criterion, or alternatively, exclude connections based on the FC connectome rather than the SC connectome (for example, using partial correlation methods that can isolate the independent contribution of direct links to FC, see ref. 36, 37). Lastly, future studies should try to address the above intrinsic structural-functional discrepancy after all. Though challenging, it has the premise to make multimodal techniques, as the one presented here, greater both in scope and in accuracy.

## Materials and methods

### Data acquisition and processing

For this study, we used the “minimally pre-processed” diffusion and functional MRI data of 50 healthy young adult subjects from the Human Connectome Project (HCP) (38). The MRI data were acquired on a customized Siemens Magnetom Skyra 3T MRI system using a multi-band pulse sequence (39–45), and included a rigorous pre-processing pipeline (44), thus minimising many of the measurement errors (see *SI appendix* for detailed acquisition and pre-processing methods used in the Human Connectome Project). We then applied further processing in order to generate functional and structural connectomes (*SI appendix*), utilising pipelines that minimise the biases introduced by acquisition, as well as processing, of both diffusion MRI (9, 46) and resting-state fMRI (13, 47). Due to its importance to the validity of the metric, our approach to minimising tractography biases is also summarised below.

### Subject-level minimisation of tractography biases in SC

To optimise the subsequent minimisation of tractography biases, the generation of tractogram *Tk* for each subject *k* included multi-shell multi-tissue constrained spherical-deconvolution (48–50) followed by probabilistic tractography of 10 million streamlines (51) with anatomically-constrained tractography (ACT, ref. 52). We then applied the SIFT2 technique separately for each individual (15). The SIFT family of algorithms reduces biases in tractography by relating local streamlines densities to the diffusion signal used for their reconstruction (15, 19). It was shown that SIFT minimises within- and between-subject measurement errors of SC (34), and as a whole, makes SC more biologically meaningful. In order to reduce the biases, SIFT2 does not rely on streamline count, a method which has been repeatedly shown to be inaccurate (9, 46, 53). Instead, it calculates connection strength as a weighted sum of streamlines, with the weights of the streamlines being estimated directly from the processed diffusion signal *S*_*k*_. The above procedure can be expressed as:

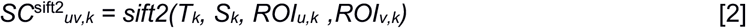

with *SC*^sift2^_*uv,k*_ being the structural connection strength between brain regions *u* and *v*, and *ROI*_*u,k*_ and *ROI*_*v,k*_ are regions-of-interest defined in subject space. Processing of the structural data and visualisation were performed with *MRtrix3* (www.mrtrix.org).

Other than the general biases in SC and FC processing, there are also challenges specific to the joint analysis of the modalities. Below we describe the steps that should be taken to address them before the FC-SC mismatch metric can be used meaningfully.

### Data-driven transformation of SC

This step addresses the discrepancy between the highly non-normal distribution of SC (20) and the approximately normal distribution of FC; a situation that invalidates the homoscedasticity assumption when using linear regression models to predict one modality from the other. Instead of resorting to complex non-linear predictive models (e.g., ref. 5), the calculation of the FC-SC metric incorporates a power-law transformation of SC (cf. ref. 35) that results in the distribution of SC becoming similar to that of FC. Because both distributions are then close to a normal distribution, the correlation between SC and FC is also expected to improve (3).

Our and others’ choice in a power-law transformation is motivated by the prevalence of power-law distributions in the neural system (54). However, in contrast with a previous study that depended on arbitrary parameters/heuristics for the power-law (35), we favour a data-driven approach. As the goal is to match the *distributions* (rather than the values) of SC and FC, the first step is to rank both SC and FC in ascending order. Then, it is possible to find the power-law parameters *a*, *b* and *c* (see Eq. 3 below) whose application to the ordered SC will best-match each SC value with its equal-rank FC value. This mathematical operation is equivalent to fitting a power-law function to the curve defined by ordered SC (x-axis) and ordered FC (y-axis). Here, we fit the curve by minimizing the least absolute residuals, using the Levenberg-Marquardt algorithm (Matlab’s curve-fitting toolbox, MathWorks, Natick, MA).

For the sake of numerical stability, the *a*, *b* and *c* parameters are estimated from the group-averaged SC and FC rather than from individual SC and FC. They are then applied to each individual separately:

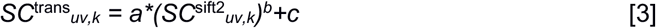

It is of note that although previous studies manipulated the distribution of SC as well, this was usually done by resampling SC (only preserving rank) into a specific target distribution, e.g., Gaussian (3), rather than using a data-driven approach like here. We opted against the resampling approach as it discards the proportional relationship between connection strengths, and could potentially reduce large differences between left and right homologues or introduce differences where such do not exist.

### Graph theory-based connection exclusion

This step addresses another intrinsic discrepancy between the modalities: the fact that SC only describes direct links, whereas FC is potentially mediated by both direct and indirect links. To be able to integrate SC and FC, one of these modalities might be altered to have the same scope as its companion. Unfortunately, however, including in SC all indirect links is intractable (because the SC connectome is an almost fully-connected weighted graph, see ref. 20, tracing all possible indirect links is a factorial problem), and eliminating from FC the contribution of such links is mathematically ill-posed, and thus, can be only approximate (e.g., ref. 12). Instead of exploring solutions to bridge this gap (4), we decided to simply exclude from analysis all those connections where indirect links have major contribution to connectivity. It limits the scope of the FC-SC metric, but at the same time, increases its interpretability.

Our criterion for excluding a connection between regions (or ‘nodes’ in the graph) *u* and *v* is the existence of an indirect path *PI*_*uv*_ = {*SC*^trans^_*ur*_, *SC*^trans^_*rs*_, …, *SC*^trans^_*tv*_} that is shorter than the direct path *PDuv* = {*SC*^trans^_*uv*_} between the two regions (a ‘path’ in the graph represents a structural link; it traverses one or more ‘edges’ of the graph). Notice that in graph theory, the length of a path in weighted graphs does not correspond to the number of edges that constitute it, and that is why the direct path is not automatically also the shortest. Instead, path length is assumed to quantify the ease of information transfer through the path, and for a graph that describes brain connectivity like here, a shorter indirect path indicates higher physiological efficacy in indirect links. This means that, on the whole, *FC*_*uv*_ is mainly driven by indirect links (be it one or more of them), and based on the criterion presented above, connection {*u*,*v*} should be excluded. For the calculation of the metric, we define the complimentary set, i.e. those connections that will not be excluded, as all connections where the direct path is the shortest of all paths:

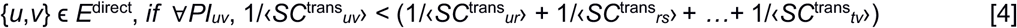

Notice that the length of the indirect path in the formula is the sum of the reciprocals of the edges that constitute it (a popular convention). Consequently, an edge with strong structural connectivity (e.g., thick fibre bundle) adds to the sum only a small amount, which contributes towards a shorter path length, i.e. higher efficacy. Lastly, because this study compares between the hemispheres, *E*^direct^ was further restricted to only these intra-hemispheric connections where the above formula is true bilaterally.

This procedure may arguably exclude a connection {*u,v*} even if the two regions are not communicating through any indirect links, and the short indirect path between them is formed by a third region that sends strong projections to both. This behaviour of the method is not necessarily negative, though, as taking into account the joint excitation of two regions by a shared external influence is yet another hurdle for models that predict FC from SC (55). As there are no practical solutions for this difficulty, excluding affected connections from analysis may be the lesser of two evils.

### Exclusion of inter-hemispheric connections

The FC-SC relationship is known to vary between intra-hemispheric and inter-hemispheric connections (see ref. 56 and Fig. 1*D*). In order to exclude this source of variability, we recommend on calculating the metric for either intra-hemispheric connections, *E*^intra^ (this study), or inter-hemispheric connections, but not both in the same analysis.

It is of note that the proposed exclusions do not only limit the scope of the metric, but also limit the set of connections used for estimating the predictive linear model between SC and FC (see *Metric*). This improves the model, as it is now not influenced by biases due to indirect links or mixing intra- and inter-hemispheric connections. Lastly, although the exclusions can be equally applied to each individual separately, we performed the exclusion at group-level (see Fig. 1). This prevents situations where connections are excluded from only some of the subjects, which may complicate statistical analysis.

## Acknowledgements

We are grateful for the support of the National Health and Medical Research Council of Australia (grant numbers APP1091593 and APP1117724); the Australian Research Council (grant number DP170101815); the Victorian Government’s Operational Infrastructure Support; and of Melbourne Bioinformatics at the University of Melbourne (grant number UOM0048). OC is supported by fellowship funding from the National Imaging Facility (NIF), a National Collaborative Research Infrastructure Strategy (NCRIS) capability at Swinburne Neuroimaging, Swinburne University of Technology. We also acknowledge the Sydney Informatics Hub and the University of Sydney’s high performance computing cluster Artemis for providing some of the high performance computing resources that have contributed to the research results reported within this paper. Data were provided by the Human Connectome Project, WU-Minn Consortium (Principal Investigators: David Van Essen and Kamil Ugurbil; 1U54MH091657) funded by the 16 NIH Institutes and Centers that support the NIH Blueprint for Neuroscience Research; and by the McDonnell Center for Systems Neuroscience at Washington University, St. Louis, MO. Lastly, we would like to thank Xiaoyun Liang for his assistance and suggestions.

