## Supplementary Information for "A functional-structural connectivity metric detects ipsilateral connections with distinct functional specialisation in each hemisphere"

Oren Civier, Marion Sourty, Fernando Calamante

Oren Civier


### Supplementary Information Text

We analysed 50 pre-processed adult datasets from the Human Connectome Project (HCP) (1), which were acquired on a customized Siemens Magnetom Skyra 3T MRI system using a multi-band pulse sequence (2-5).

#### Structural data - grey matter

The HCP data include high resolution T1 anatomical images, acquired using the 3D magnetization-prepared rapid gradient echo sequence (MPRAGE) (6) with 0.7 x 0.7 x 0.7 $mm^3$ voxel size, TR/TE = 2400/2.14 ms, and flip angle = 8° (7, 8).

Using these data, we parcellated the brain into the 84 regions-of-interest of the Desikan-Kiliany atlas (9). Cortex and cerebellum were parcellated using freesurfer (9), whereas subcortical parcellation relied on FIRST (10, see ref. 11).

#### Structural data – white matter

**Acquisition and pre-processing.** The diffusion imaging protocol consisted of 3 diffusion-weighted shells (b = 1000, 2000, and 3000 s/$mm^2$), with 90 diffusion-weighted directions each, and 18 reference volumes (b = 0 s/$mm^2$). For distortion correction, all images were additionally acquired with reversed phase encoding (12). Other diffusion-weighted imaging parameters were: 145 x 145 matrix, 174 slices, 1.25 x 1.25 x 1.25 $mm^3$ voxel size, TR/TE = 5520/89.5 ms. The HCP pre-processing pipeline incorporates image reconstruction with SENSE1 multi-channel (13), and diffusion imaging distortion correction (14, 15).

To further process the "minimally pre-processed" HCP data consisting of vowel-wise diffusion weighted measurements, we first applied bias-field correction (16). This was followed by multi-shell multi-tissue constrained spherical deconvolution (CSD) (17, 18) to model white matter, grey matter and cerebrospinal fluid (19), with a maximum spherical harmonic degree $L_{max}$ = 8.

**Connectome generation.** For each subject, tractogram construction included several steps: generation of 10 million probabilistic streamlines using the 2$^{nd}$-order Integration over Fibre Orientation Distributions algorithm (iFOD2, see ref. 20) and anatomically-constrained tractography (ACT) (21), with dynamic seeding (22): FOD amplitude threshold 0.06, step size was half of voxel size, length of 5-300 mm, and backtracking, i.e. possibility for tracks to be truncated and re-tracked if a poor structural termination is encountered (21). In the next step, each streamline was assigned a weight computed using SIFT2 (22); to achieve that, SIFT2 runs global optimisation to select weights that would fit the tractogram to the underlying data best. Connection strengths were calculated by summing the weights of the streamlines that connect each pair of parcellated regions-of-interest.

#### Functional data

**Acquisition and pre-processing.** The resting-state functional MRI protocol of this dataset used the following parameters: TR = 720 ms, using a multiband factor of 8; TE = 33.1 ms; flip angle = 52°, 2 mm isotropic resolution (FOV: 208 mm × 180 mm, Matrix: 104 × 90 with 72 slices). The four available runs of 14 min and 33 sec each (1200 volumes) were analysed separately in this study (7, 8).

Pre-processing steps are described in detail elsewhere (8). Briefly, pre-processing included gradient distortion correction, motion correction, fieldmap-based EPI distortion correction, brain-boundary-based registration of EPI to structural T1-weighted volume, non-linear registration into MNIspace, and intensity normalization. The images are then denoised using the ICA-FIX method (FMRIB's ICA-based Xnoiseifier) (23, 24).

**Connectome generation.** The pre-processed data consisted of voxel-wise time series representing denoised BOLD signal fluctuations. We analysed these data separately for each subject *k*. Using the same 84-nodes parcellation as previously described, we calculated the time-course of each region-of-interest by averaging the BOLD signals of its voxels, separately for each of the four runs. The functional connectome of each run was computed using Pearson's correlation coefficient between the time-course of each pair of regions-of-interest. For each subject, the four runs are then averaged to form a single mean functional connectome. This averaged out run-specific effects (range of cognitive and attentional states, influence of head motion, quality of registration to structural scan, etc.), reducing inter-subject variability in FC (25, 26). Processing of the functional data was performed with Matlab (MathWorks, Natick, MA).

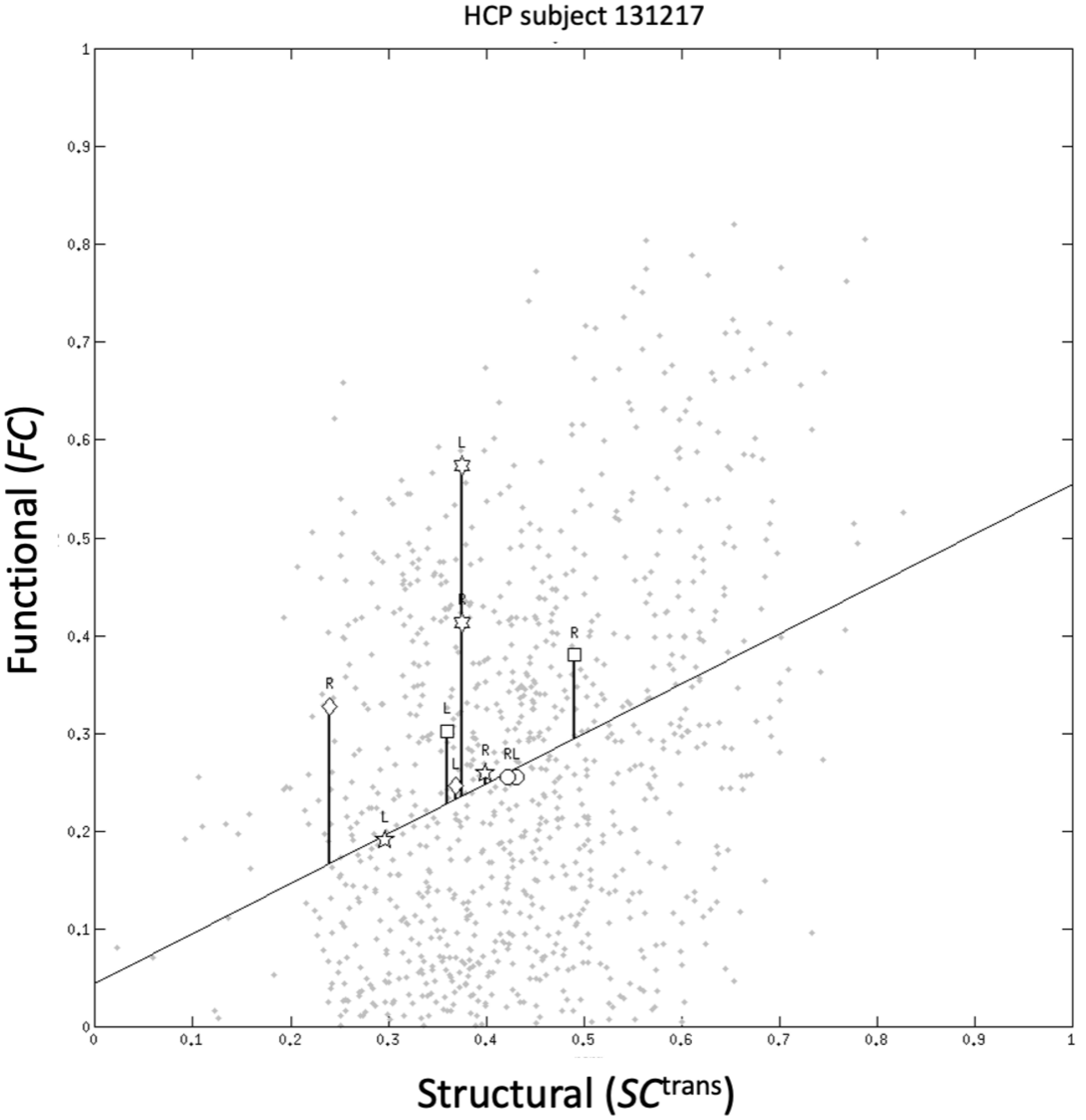


**Figure S1**. Scatterplot of empirical *FC* against $SC^{trans}$ (grey points) and corresponding regression line for a representative subject from the Human Connectome Project (HCP) dataset. The y-axis distance of each data point from the regression line is equal to its FC-SC mismatch metric ($mFCSC_{uv,k}$). The data points for several bilateral pairs are indicated: Circles - pars operculis-inferior parietal, Pentagrams - inferior temporal-banks of superior temporal sulcus, Squares - rostral middle frontal-superior parietal, Hexagrams - superior frontal-posterior cingulate, Diamonds - caudal middle frontal-lateral occipital (notice that a pattern at the level of a single subject might not reflect the common pattern across the group). Conventions are as in Fig. 2 of the main text.

**Table S1.** Bilateral pairs where there is a significant difference between the FC-SC mismatch metric of the left and right homologues.

| The relationship between the left FC-SC mismatch metric value, right FC-SC mismatch metric value, and zero | Bilateral pairs of intra-hemispheric connections[a] |
|---|---|
| (a) $\langle mFCSC_{[L]} \rangle > \langle mFCSC_{[R]} \rangle > 0$ | superior frontal-pars opercularis[b]<br>banks of the superior temporal sulcus-caudal middle frontal<br>caudal middle frontal-inferior temporal<br>inferior parietal-pars opercularis<br>inferior temporal-pars orbitalis<br>pars opercularis-pars orbitalis<br>caudal middle frontal-pars triangularis<br>inferior parietal-pars triangularis<br>middle temporal-pars triangularis<br>pars opercularis-pars triangularis<br>pars orbitalis-pars triangularis<br>lingual-pericalcarine<br>isthmus cingulate-rostral anterior cingulate<br>inferior temporal-rostral middle frontal<br>rostral middle frontal-superior parietal<br>lateral orbitofrontal-superior temporal<br>rostral middle frontal-supramarginal |
| (b) $\langle mFCSC_{[L]} \rangle > 0 > \langle mFCSC_{[R]} \rangle$ | middle temporal-pars opercularis<br>postcentral-rostral middle frontal<br>precentral-rostral middle frontal<br>medial orbitofrontal-superior frontal<br>caudal anterior cingulate-thalamus |
| (c) $0 > \langle mFCSC_{[L]} \rangle > \langle mFCSC_{[R]} \rangle$ | medial orbitofrontal-pars orbitalis<br>pars orbitalis-precentral<br>lateral orbitofrontal-rostral anterior cingulate |
| (d) $\langle mFCSC_{[L]} \rangle < \langle mFCSC_{[R]} \rangle < 0$ | pars opercularis-putamen<br>pars triangularis-putamen |
| (e) $\langle mFCSC_{[L]} \rangle < 0 < \langle mFCSC_{[R]} \rangle$ | caudal middle frontal-insula<br>inferior parietal-insula<br>pars triangularis-insula<br>postcentral-hippocampus |
| (f) $0 < \langle mFCSC_{[L]} \rangle < \langle mFCSC_{[R]} \rangle$ | banks of the superior temporal sulcus-precentral<br>precentral-superior temporal<br>precentral-supramarginal<br>banks of the superior temporal sulcus-insula<br>pars opercularis-insula<br>precentral-insula<br>precuneus-insula<br>superior frontal-insula<br>supramarginal-insula<br>superior frontal-cerebellum |

[a] Bilateral pairs are named after the two brain regions that define their intra-hemispheric connections. The order of the two regions is arbitrary, as each intra-hemispheric connection may include axons that project either way.
[b] Bilateral pair is shown in Fig. 2 of the main text for selected subjects.

**Table S2.** Bilateral pairs with significant functional connectivity asymmetry, and where the asymmetry reflects the direction and extent of hemispheric dominance.

| Functional Connectivity Asymmetry | Bilateral pairs of intra-hemispheric connections[a] |
|---|---|
| (a) $FC_{[L]} > FC_{[R]}$ reflecting the extent of a leftward dominance | banks of the superior temporal sulcus-middle temporal<br>banks of the superior temporal sulcus-pars opercularis<br>caudal middle frontal-pars opercularis<br>medial orbitofrontal-rostral anterior cingulate<br>posterior cingulate-rostral anterior cingulate<br>precuneus-rostral anterior cingulate<br>fusiform-rostral middle frontal<br>caudal middle frontal-superior temporal<br>pars opercularis-superior temporal<br>middle temporal-supramarginal<br>supramarginal-cerebellum |
| (b) $FC_{[L]} < FC_{[R]}$ reflecting the extent of a rightward dominance | inferior parietal-rostral middle frontal |

[a] See footnote a in Table S1.
